# CRACD suppresses neuroendocrinal plasticity of lung adenocarcinoma

**DOI:** 10.1101/2023.04.19.537576

**Authors:** Bongjun Kim, Shengzhe Zhang, Yuanjian Huang, Kyung-Pil Ko, Gengyi Zou, Jie Zhang, Sohee Jun, Kee-Beom Kim, Youn-Sang Jung, Kwon-Sik Park, Jae-Il Park

## Abstract

Tumor cell plasticity contributes to intratumoral heterogeneity and therapy resistance. Through cell plasticity, lung adenocarcinoma (LUAD) cells transform into neuroendocrinal (NE) tumor cells. However, the mechanisms of NE cell plasticity remain unclear. CRACD, a capping protein inhibitor, is frequently inactivated in cancers. *CRACD* knock-out (KO) de-represses NE-related gene expression in the pulmonary epithelium and LUAD cells. In LUAD mouse models, *Cracd* KO increases intratumoral heterogeneity with NE gene expression. Single-cell transcriptomic analysis showed that *Cracd* KO-induced NE plasticity is associated with cell de-differentiation and activated stemness-related pathways. The single-cell transcriptomes of LUAD patient tumors recapitulate that the distinct LUAD NE cell cluster expressing NE genes is co-enriched with SOX2, OCT4, and NANOG pathway activation, and impaired actin remodeling. This study reveals an unexpected role of CRACD in restricting NE cell plasticity that induces cell de-differentiation, providing new insights into cell plasticity of LUAD.

## Introduction

Cell plasticity, a process changing cell fate or state ^1–3^, plays pivotal roles in development, tissue homeostasis, and regeneration. During development, embryonic progenitor cells change their cell fate ^2, 3^. Upon cell intrinsic or extrinsic signaling cues, terminally differentiated cells undergo cell plasticity via de-differentiation or trans-differentiation, contributing to homeostasis and regeneration of many tissues ^4–10^.

Cell plasticity plays also a crucial role in tumorigenesis ^11, 12^. Tumor cell plasticity is associated with tumor progression, intratumoral heterogeneity, and therapy resistance ^11–13^. In LUAD, tumor cell plasticity changes the cancer subtype ^12, 14, 15^. For example, during EGFR targeted therapies, *EGFR* mutant LUAD tumor cells transform into NE tumor cells ^16, 17^. A Kras^G12C^ inhibitor, AMG510, induces tumor cell plasticity converting *KRAS*^G12C^ mutant LUAD tumor cells into squamous cancer cells ^18^. The ALK inhibitor, crizotinib, changes *ALK*-mutant LUAD tumor cells into small cell lung cancer (SCLC) ^19^. NE cell plasticity was also observed in melanoma ^20^, pancreatic adenocarcinoma ^21^ and prostate cancer ^22^. However, the mechanisms of NE cell plasticity of LUAD remain elusive.

In this study, leveraging genetically engineered mouse models, organoids, and single-cell transcriptomics, we found that CRACD tumor suppressor serves as a gatekeeper restricting NE cell plasticity, which might be implicated in LUAD’s therapy resistance and tumor cell heterogeneity.

## Results

### *Cracd* KO generates NE-like pulmonary epithelial cells

Previously, we identified the CRACD (<u>C</u>apping protein inhibiting <u>R</u>egulator of <u>Ac</u>tin Dynamics; also known as CRAD/KIAA1211) tumor suppressor, which promotes actin polymerization by binding and inhibiting capping proteins to promote actin polymerization ^23^. Interestingly, we observed SCLC-like lesions in the lungs of *Cracd* KO mice ^23^. This observation led us to hypothesize that CRACD loss may drive NE-like cell plasticity in the lung. To test this, we examined *Cracd* KO mouse lung tissues. Unlike *Cracd* wild-type (WT), *Cracd* KO lung tissues showed NE-like hyperplasia in the bronchiolar airway and alveoli (Fig. 1A). Immunofluorescent (IF) staining confirmed the proliferative nature of this NE-like cell mass, as indicated by MKI67+ IF staining. Furthermore, the mass expressed several NE markers, including KRT19, SYP, CGRP, and CHGA (Fig. 1B). It is noteworthy that *Cracd* KO alone failed to develop lung tumors in mice ^23^. We also assessed the expression of NE markers in lung organoids (LOs) derived from pulmonary epithelial cells isolated from murine lung tissues (*Cracd* WT vs. KO) ^24^ (Fig. 1C, D; fig. S1). We confirmed the generation of three different types of LOs: alveolar (HOPX+, SPC+), bronchiolar (Ac-TUB+, SCGB1A1+), and bronchioalveolar (HOPX+, SPC+, Ac-TUB+, SCGB1A1+) types (Fig. 1E). The *Cracd* KO LOs exhibited increased expression of NE markers, CHGA and CGRP, in both bronchiolar and alveolar LOs (Fig. 1F, G). These results suggest that CRACD loss is sufficient to induce the expression of NE-like features in the pulmonary epithelium.

**Figure 1.**
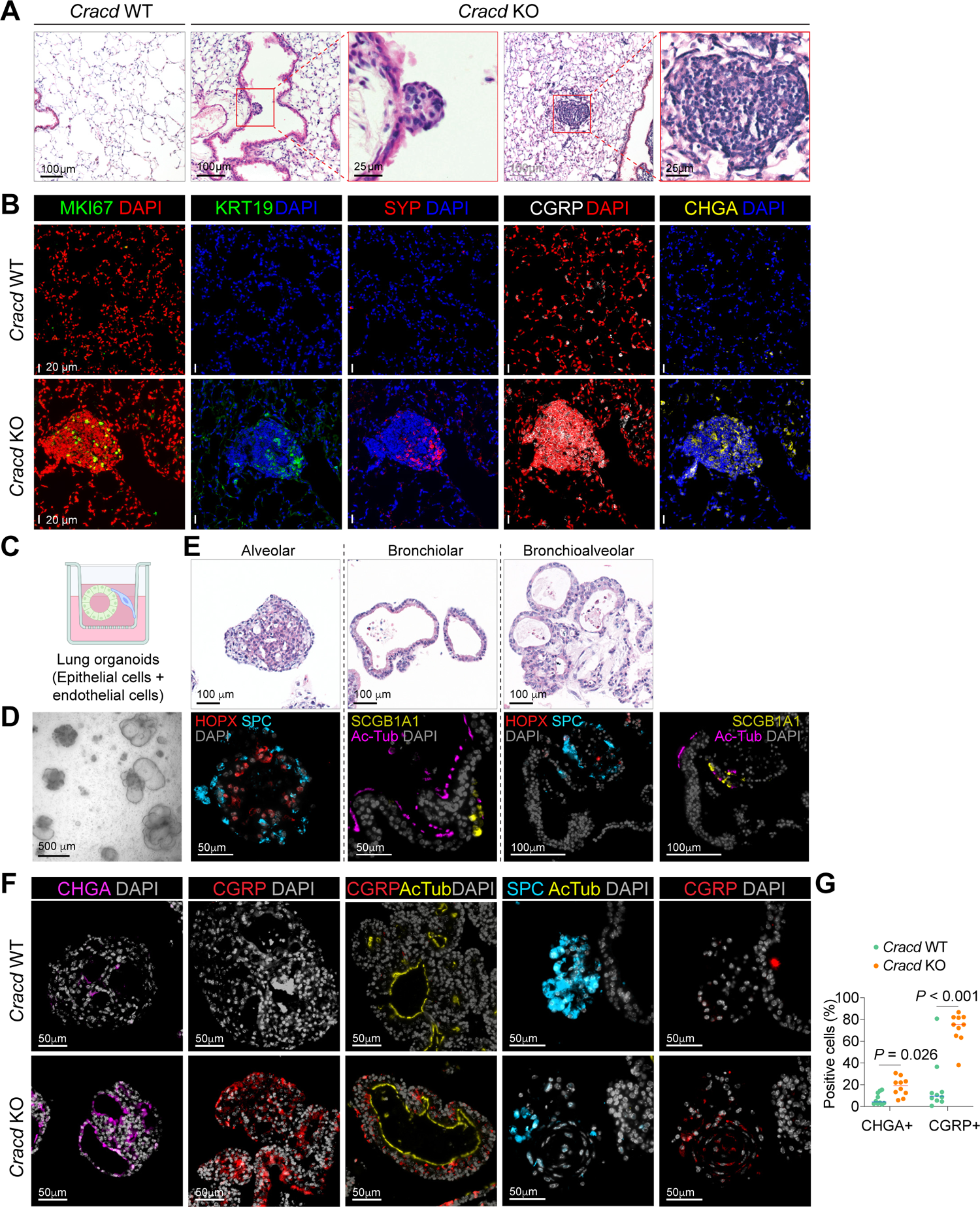
*Cracd* KO induces NE cell-like features in the pulmonary epithelium and organoids. **A, B,** Hematoxylin and eosin (H&E) (A) and immunofluorescent (IF) (B) staining of mouse lung sections (*Cracd* WT vs. KO) mice (n = 3 per group). **C,** Illustration of lung organoid culture. **D,** Bright-field images of lung organoids (LOs) at day 12. **E,** H&E (upper panels) and IF (lower panels) staining of LOs. **F,** IF staining of LOs derived from *Cracd* WT vs. KO mice. **G,** Quantification of CHGA+ and CGRP+ cells in LOs (n = 10 per LO). Two-tailed Student’s *t*-test; error bars: SD. Representative images were displayed.

### CRACD depletion upregulates NE marker genes in LUAD cells

Having observed NE-like features in *Cracd* KO lung, we investigated whether CRACD depletion also induces NE marker expression in non-NE tumor cells, particularly LUAD cells. We introduced CRACD shRNA into both mouse (KP-1, derived from *<u>K</u>ras*^G12D^; *Tr<u>p</u>53* KO mouse LUAD tumors) ^25^ and human (A549) LUAD cell lines. We found that CRACD depletion upregulated the expression of NE marker genes in both KP-1 and A549 cells, compared to control cells (Fig. 2A). Moreover, CRACD depletion led to a reduction in the cytoplasmic-to-nuclear ratio with the loss of F-actin stress fibers (Fig. 2B, C), confirming the role of CRACD in maintaining actin polymerization.

**Figure 2.**
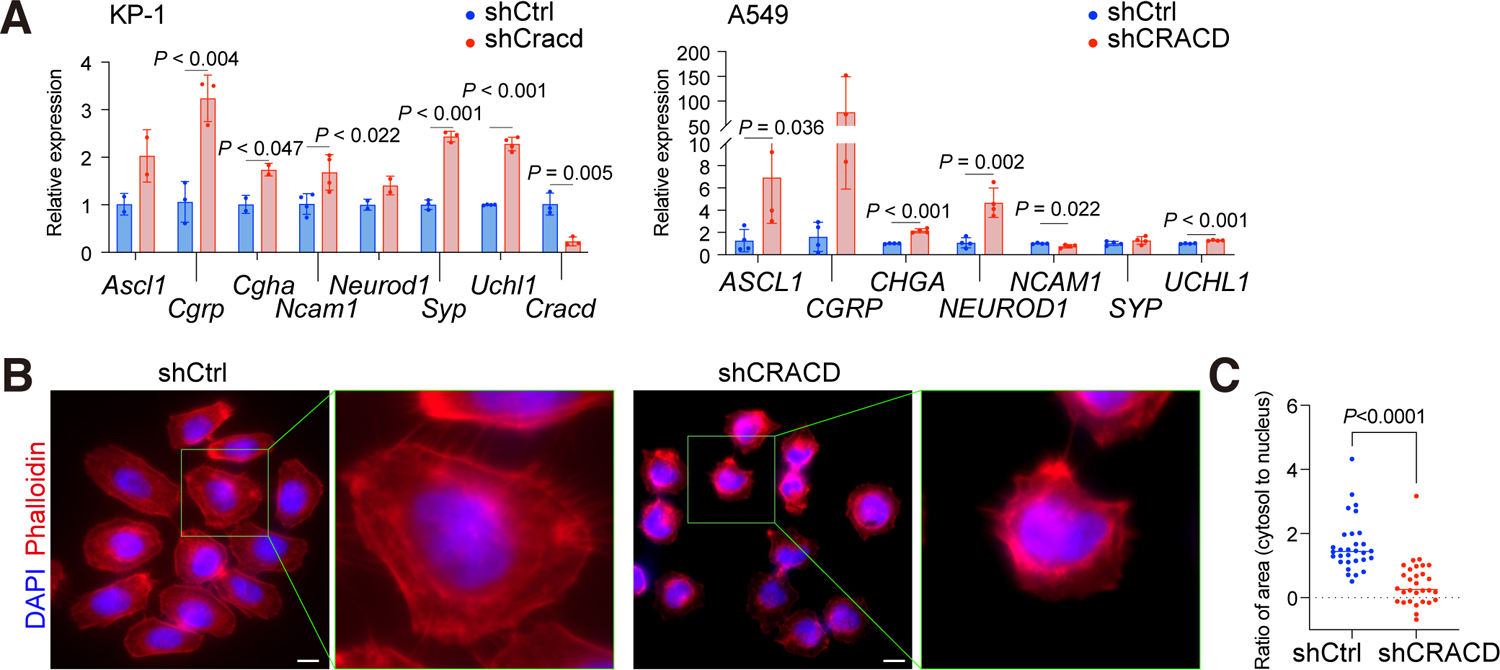
CRACD depletion de-represses NE gene expression in LUAD cells. **A,** qRT-PCR analysis of KP-1 cells (left panel) and A549 cells (right panel) stably transduced with the lentiviruses encoding *shCracd* or *shCRACD*, respectively; two-tailed Student’s *t*-test; error bars: SD. **B,** IF staining of A549 cells (*shCtrl* vs. *shCRACD*) for phalloidin, a marker for filamentous actin (n = 3). Representative images were shown. **C,** Quantification of cytosol-to-nucleus ratio of images (Fig. 2B) (n = 30).

### *Cracd* KO induces NE cell plasticity in LUAD driven by *Kras*^G12D^ and *Trp53* KO

Next, we determined the impact of CRACD loss on the plasticity of LUAD tumor cells in vivo. To genetically ablate *Cracd* alleles in vivo, we employed two approaches: CRISPR-based somatic gene targeting ^26^ and germline deletion. For somatic engineering, we administered adenovirus harboring Cas9-sgLacZ-Cre (control) or Cas9-sgCracd-Cre into KP (*<u>K</u>ras*^G12D/WT^; *Tr<u>p</u>53* ^f/f^ ^(floxed/floxed)^) mice, a LUAD mice model, via intratracheal instillation (Fig. 3A). Twelve weeks after adenovirus treatment, we collected lung tissues for tumor analyses. Compared to *Cracd* WT KP-induced LUAD (control), *Cracd* KO KP tumors exhibited significant heterogeneity in tumor cell morphology (Fig. 3B, C, fig. S2). Moreover, unlike *Cracd* WT KP LUAD where NE markers were rarely expressed, *Cracd* KO KP tumors showed the expression of NE markers, such as CHGA, CGRP, and ASCL1 (Fig. 3D). We confirmed that the NE-marker expressing *Cracd* KO KP cells are tumor cells by performing CDH1/E-cadherin IF staining (Fig. 3E). Additionally, *Cracd* KO tumor cells showed disrupted actin cytoskeleton (Fig. 3E). To complement the somatic engineering, we also established the *Cracd* KO (heterozygous and homozygous); *<u>K</u>ras*^G12D^; *Tr<u>p</u>53*^f/f^ (CKP) compound strain. To induce LUAD development, we administered Cre recombinase-expressing adenovirus (Ad-Cre) to KP (control) and CKP mice via intratracheal instillation. Twelve weeks after administration, we collected lung tumors for analyses (Fig. 3F). Consistent with the results of somatic engineering, KP tumors carrying the germline mutation of *Cracd* exhibited marked expression of CHGA, CGRP, and NEUROD1, and disrupted actin structure, while *Cracd* WT KP tumors did not (Fig. 3G, H). Moreover, both *Cracd* homozygous KO (-/-) and heterozygous (+/-) tumors showed increased intratumoral heterogeneity (Fig. 3I, J). These results suggest that CRACD loss is sufficient to de-repress NE-related genes and increase intratumoral heterogeneity in LUAD.

**Figure 3.**
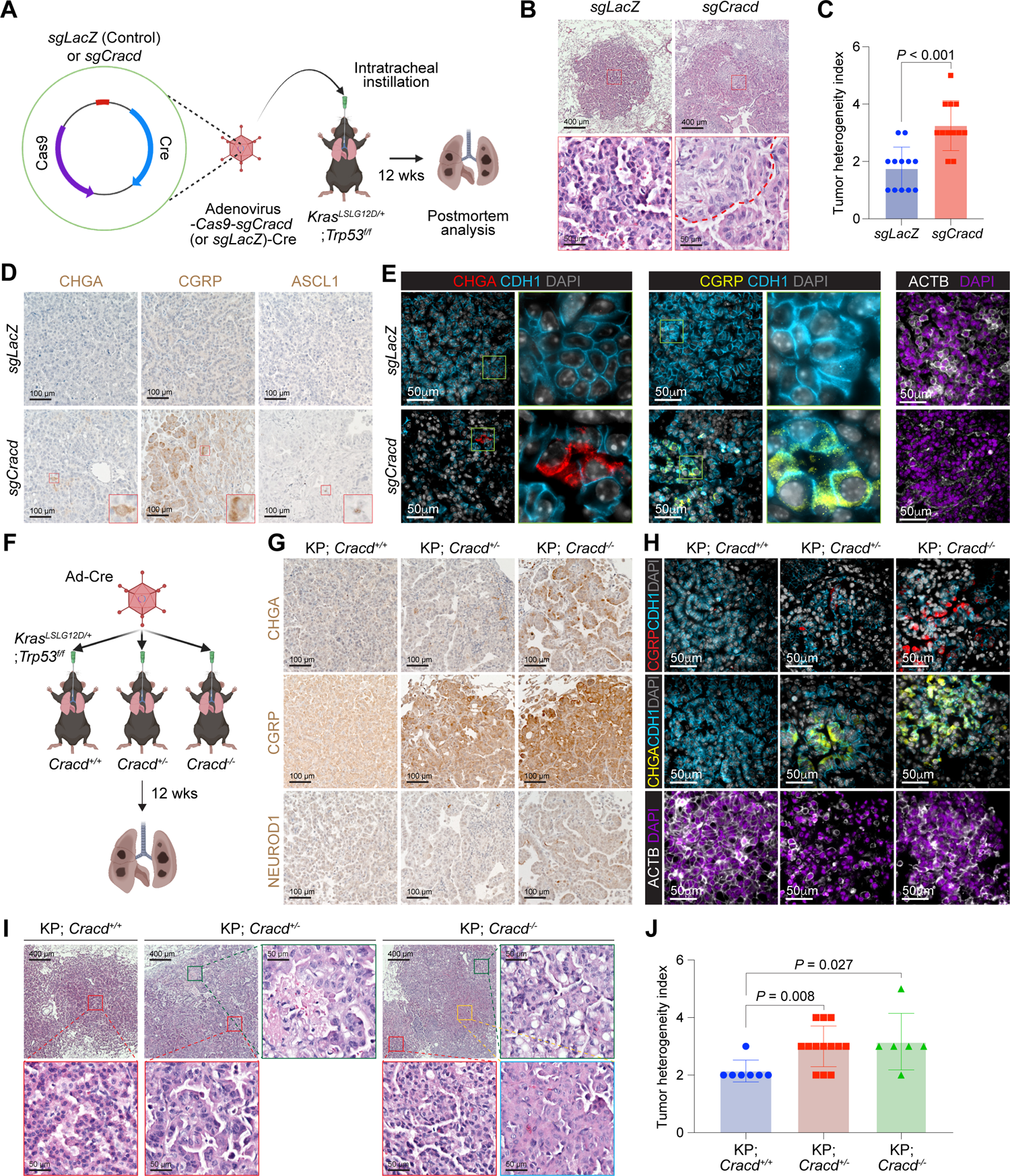
*Cracd* KO increases tumor heterogeneity with NE gene expression in LUAD mouse models. **A,** Illustration of somatic gene targeting using adenovirus encoding sgRNAs and Cre; (n = 3 per group). **B, C,** Tumor heterogeneity analysis; H&E (B); intratumoral heterogeneity index (C) (n = 12 per group). **D, E,** Immunostaining of lung tumors; DAB (3,3’-Diaminobenzidine) (D); IF (E). **F,** Experimental scheme of Cracd-deficient LUAD mice model using Cracd germline KO mice. **G,H,** Immunostaining of lung tumors; DAB (G); IF (H). **I, J,** Tumor heterogeneity analysis; H&E (I); intratumoral heterogeneity index (J); WT (n = 3) vs. heterozygous (n=11) vs. homozygous (n=2). Representative images were shown. Two-tailed Student’s *t*-test; error bars: SD.

### NE cell plasticity is associated with cell de-differentiation of pulmonary epithelial and LUAD cells

To elucidate the mechanisms of *Cracd* KO-induced NE marker expression and cellular heterogeneity increase in LUAD, we employed single-cell transcriptomics. We isolated pulmonary epithelial cells from mouse lung tissues (*Cracd* WT or KO) and performed single-cell RNA sequencing (scRNA-seq) and comparative analyses (fig. S3). Using unsupervised clustering and annotations, we identified each pulmonary epithelial cell type (Fig. 4A, B; fig. S3, Table S2). Consistent with the IF results (Fig. 1), the *Cracd* KO lung tissue exhibited relatively higher expression of NE- and SCLC-related genes (Fig. 4C). Since cell plasticity is associated with cell de-differentiation or transdifferentiation, we evaluated the impact of *Cracd* KO on cell differentiation and de-differentiation states, we used the CytoTRACE package that infers cell differentiation state by RNA content ^27^. Notably, the *Cracd* KO AT2 clusters (AT2-1∼6 cell clusters) displayed significantly less-differentiated states compared to those of *Cracd* WT (Fig. 4D). To determine the signaling pathways involved in *Cracd* KO-induced NE cell plasticity, we conducted fGSEA (fast Geneset Enrichment Analysis) and found that cell stemness-related gene signatures, including OCT4, and NANOG targets (Table S3) ^28^, were highly enriched in the AT2 cell clusters of the *Cracd* KO lung tissues compared to WT (Fig. 4E), which was shown by the dot and feature plots (Fig. 4F, G).

**Figure 4.**
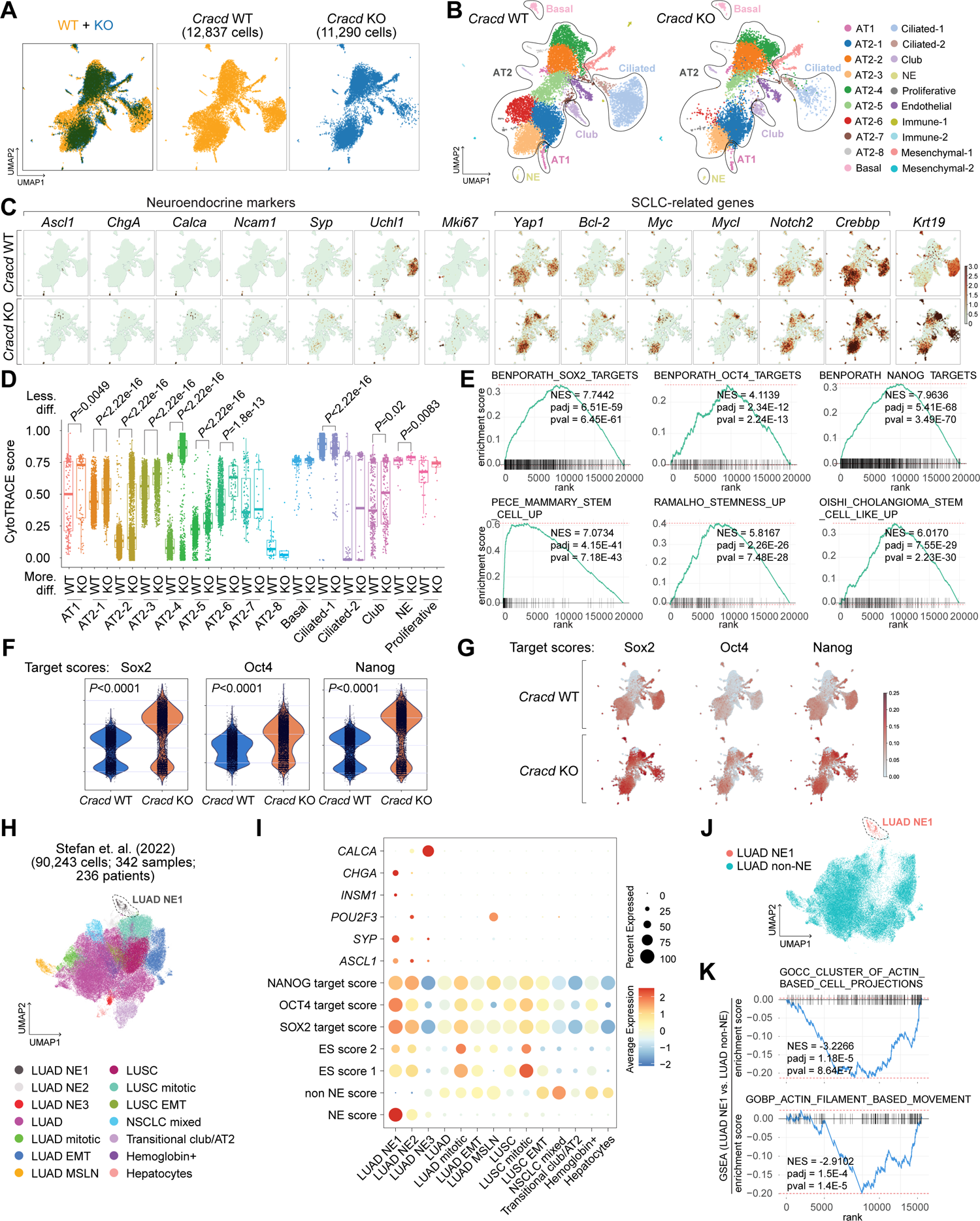
Association of NE cell plasticity with cell stemness in pulmonary epithelial and LUAD tumor cells. **A,** Uniform manifold approximation and projection (UMAP) plots displaying pulmonary epithelial cells from *Cracd* WT vs. KO mice. **B,** UMAPs of each cell cluster annotated by cell types. **C,** Feature plots showing the expression of NE- or SCLC-related genes. **D,** Boxplots of CytoTRACE scores of each cell cluster; less/more diff: less/more differentiated cell states. **E,** GSEA of the AT2 clusters (*Cracd* WT vs. KO) using the datasets shown in Figure 4A. **F,** Dot plots depicting transcriptional module scores of Sox2, Oct4, and Nanog in AT2 clusters. **G,** Feature plots showing the module scores (Sox2, Oct4, and Nanog). **H,** UMAP of NSCLC tumor cells annotated by tumor cell types. **I,** Dot plot depicting NE gene expression and transcriptional module scores of the gene sets. **J,** UMAP displaying the two subsets (LUAD NE1 vs. LUAD non-NE [LUAD, LUAD mitotic, LUAD EMT, and LUAD MSLN]). **K,** GSEA of LUAD NE1 vs. LUAD non-NE.

Subsequently, to assess the pathological relevance of the association between NE cell plasticity and cell de-differentiation in LUAD, we analyzed scRNA-seq datasets of non-small cell lung cancer (NSCLC) patient tumor samples ^29^. We re-analyze the pre-processed dataset of epithelial compartments consisting of 342 datasets (90,243 tumor cells), and refined the clusters into different types of tumor cells, including LUAD (mitotic, EMT, MSLN [*MSLN* high]), LUAD NE1-3 (neuroendocrine), and lung squamous cell cancer (LUSC) cells (mitotic and EMT) (Fig. 4H), as previously described ^29^. We then determined whether NE-related genes were co-expressed with stemness-related genes in LUAD. Among all clusters, the LUAD NE1 cell cluster exhibited high NE score, including NE-related genes (*CHGA*, *INSM1*, *SYP*, and *ASCL1*), and stemness-related genes (target genes of SOX2, OCT4, and NANOG and stemness genes enriched in embryonic stem cells [ES]) (Fig. 4I, Table S3) ^28^, which is consistent with the results from the *Cracd* KO lung scRNA-seq analysis (Fig. 4E-G). Since CRACD loss impairs actin remodeling and induces NE cell plasticity, we asked whether the LUAD NE1 cell cluster is related to the disrupted actin pathway. Indeed, fGSEA analysis showed that actin remodeling-related pathways were decreased in the LUAD NE1 clusters compared to LUAD non-NE clusters (LUAD, LUAD mitotic, LUAD EMT, and LUAD MSLN) (Fig. 4J, K). Since CRACD inhibits the WNT signaling ^23^, we also examined the effect of *Cracd* KO on WNT signaling. The WNT pathway target genes were marginally increased in *Cracd* KO lung compared to WT (fig. S4A). Similarly, the expression of WNT signaling target genes was barely altered in LUAD NE1 clusters compared to other clusters (fig. S4B). These results suggest that NE cell plasticity is associated with cell de-differentiation of LUAD.

## Discussion

The underlying mechanisms of NE cell plasticity in LUAD are not fully understood. Genetic ablation of CRACD tumor suppressor was sufficient to de-repress NE-related genes in organoids and mice. In mice, *Cracd* KO leads to increased intratumoral heterogeneity with upregulation of NE markers. Single-cell transcriptomic analysis showed that *Cracd* KO upregulates NE-related genes primarily in AT2 pulmonary epithelial cells, accompanied by increased cell de-differentiation state. Single-cell transcriptomes of LUAD patient tumors showed the distinct LUAD NE cell cluster co-enriched with NE genes, cell stemness pathways, and impaired actin remodeling.

Tumor cell plasticity is implicated in tumor progression, intratumoral heterogeneity, and therapy resistance ^11, 12^. NE cell plasticity has been observed in lung and prostate cancer as an outcome of cancer therapy ^14, 17, 30^. Our study found that NE cell plasticity is associated with cell de-differentiation of pulmonary epithelial and LUAD tumor cells. The genetic ablation of *Cracd* alone was sufficient to induce a less differentiation state of cells (Fig. 4D). Moreover, cell stemness-related pathways were activated in *Cracd* KO pulmonary epithelial cells (Fig. 4E-G). Analysis of human LUAD single-cell transcriptomes also showed co-expression of NE and stemness-related genes (Fig. 4I). These data suggest that NE cell plasticity is likely driven or accompanied by cell de-differentiation, which implies the acquisition of cell stemness through NE cell plasticity. Cell stemness is characterized by two major features: cellular heterogeneity generation and self-renewal ^31^. Thus, such acquired cell stemness might explain why NE cell plasticity increases intratumoral heterogeneity observed in *Cracd* KO LUAD tumors (Fig. 3). Similarly, since tumor cell plasticity also contributes to therapy resistance ^11, 12^, CRACD inactivation-induced NE cell plasticity might generate therapy-resistant tumor cells. Cell plasticity is one of the hallmarks of cancer ^32^. Therefore, targeting NE cell plasticity would be an alternative option for overcoming the therapy resistance of LUAD or LUAD NE.

The *CRACD/KIAA1211* gene is frequently inactivated in SCLC ^33–36^, which somehow agrees with our finding of CRACD loss-induced NE cell plasticity since SCLC tumor cells exhibit NE features. However, the specific mechanisms by which CRACD loss-of-function takes places in LUAD remain to be determined. In colorectal cancer, CRACD inactivation occurs through transcriptional downregulation (via promoter hypermethylation) or genetic mutations (missense and nonsense). ^23^ Therefore, combined analyses of exome-seq and scRNA-seq could help determine the mechanism of CRACD inactivation in LUAD.

As a capping protein inhibitor, CRACD promotes actin polymerization. In colorectal cancer, CRACD inactivation disrupts the cadherin-catenin-actin complex, releasing β-Catenin for WNT signaling hyperactivation ^23^. Although WNT signaling was slightly activated in *Cracd* KO lung tissues (fig. S4A), WNT signaling module score in the LUAD NE cluster was barely increased (fig. S4B). Thus, it is unlikely that WNT signaling mediates CRACD inactivation-induced NE plasticity. Instead, the LUAD NE tumor cell cluster displayed relatively downregulated actin-related pathways (Fig. 4J, K). Accumulating evidence suggests that actin remodeling modulates stemness and lineage commitment ^37–39^. Therefore, it is highly probable that dysregulated actin remodeling might mediate CRACD loss-induced NE cell plasticity and increased cell de-differentiation. Mechanistically, actin cytoskeleton-driven mechanical pulling force modulates the NOTCH signaling that controls cell lineage-related genes ^40, 41^. Additionally, nuclear actin is engaged in transcriptional regulation ^42, 43^. Thus, it is possible that upon CRACD inactivation, NOTCH signaling dysregulation or epigenetic reprogramming might trigger NE cell plasticity, which needs to be addressed in future studies.

Collectively, this study reveals an unexpected role of CRACD tumor suppressor in restricting cell plasticity and cell de-differentiation, providing new insights into NE cell plasticity of LUAD.

## Acknowledgments

This work was supported by the Cancer Prevention and Research Institute of Texas (RP200315 to J.-I.P. and RP210028 to G.Z.), the National Cancer Institute (2R01 CA193297 and R03 CA256207 to J.-I.P.), and an Institutional Research Grant (MD Anderson to J.-I.P.). The core facilities at MD Anderson (DNA Sequencing and Genetically Engineered Mouse Facility) were supported by National Cancer Institute Cancer Center Support Grant (P30 CA016672). The core facilities at Baylor College of Medicine (Cytometry & Cell Sorting Core and Single Cell Genomics Core) were supported by CPRIT (RP180672, RP200504) and the National Institutes of Health (CA125123, RR024574).

## Author contributions

B.K., S.Z., and J.-I.P. conceived and designed the experiments. B.K., S.Z., Y.H., K.-P.K, G.Z., J.Z., and S.J. performed the experiments. K.-B.K. and K.-S.P. provided adenoviruses for gene targeting. B.K., S.Z., K.-S.P., and J.-I.P. analyzed the data. B.K., S.Z., and J.-I.P. wrote the manuscript.

## STAR Methods

### RESOURCE AVAILABILITY

#### Lead contact

Additional information and requests for resources and reagents should be directed to and will be fulfilled by the Lead Contact, Jae-Il Park.

#### Materials availability

The materials will be available upon request.

#### Data and code availability

scRNA-seq data are available via the Gene Expression Omnibus (GEO) and is publicly available as of the date of publication. Accession numbers are listed in the key resource table. (Log-in token for reviewers:)

R packages and python packages used in this paper are listed in the key resource table. The code used to reproduce the analyses described in this manuscript can be accessed via Zenodo (https://doi.org/) and is available upon request.

## EXPERIMENTAL MODEL AND SUBJECT DETAILS

### Mice

C57BL/6, *Trp53*^f/f^ ^(floxed/floxed)^ (JAX no. 008179), and *Kras*^G12D^ (JAX no. 008462) mice were purchased from the Jackson Laboratory. *Cracd* KO mice have been described previously ^23^. *Kras*^G12D^, *Trp53*^f/f^ ^(floxed/floxed)^ (KP), *Cracd* ^-/-^, *Kras*^G12D^, *Trp53*^f/f^ and *Cracd* ^+/-^, *Kras*^G12D^, *Trp53*^f/f^ compound strains were generated by breeding, with validation of genotypes as previously described ^23, 25^. For LUAD tumor induction, the lungs of 10-week-old mice were infected with adenoviral Cre (Ad-Cre) via intratracheal instillation as previously describe ^25, 44^. Multiple cohorts of independent litters were analyzed to control for background effects, and both male and female mice were used. For KP *sgCracd* LUAD model, adenovirus containing *sgCracd*-Cre (Ad-*sgCracd*-Cre) or *sgLacZ*-Cre (Ad-*sgLacZ*-Cre; control) were introduced into KP mice via intratracheal instillation. Ad-*sgCracd*-Cre particles were produced in Vector Development Laboratory at Baylor College of Medicine. Mice were euthanized by CO_2_ asphyxiation followed by cervical dislocation at the indicated time. Tumors were harvested from euthanized mice, fixed with 10% formalin, embedded in paraffin, and sectioned at 5-μm thickness. The sections were stained with hematoxylin and eosin for histological analysis. All mice were maintained in compliance with the guidelines of the Institutional Animal Care and Use Committee of the University of Texas MD Anderson Cancer Center. All animal procedures were performed based on the guidelines of the Association for the Assessment and Accreditation of Laboratory Animal Care and institutionally approved protocols. This study was compliant with all relevant ethical regulations regarding animal research.

### Lung cell isolation

Lung tissues were harvested from euthanized mice after perfusing 10 ml of cold phosphate-buffered saline (PBS) into the right ventricle. Lungs were minced after the removal of extra-pulmonary tissues and digested in Leibovitz media (Gibco, USA, no. 21083-027) with 2 mg/ml collagenase type I (Worthington, CLS-1, LS004197), 2 mg/ml elastase (Worthington, ESL, LS002294), and 0.4 mg/ml DNase I (Sigma, DN-25) for 45 min at 37 °C. To stop the digestion, fetal bovine serum (FBS, HyClone; Cytiva) was added to a final concentration of 20%. The digested tissues were sequentially filtered through a 70-μm and a 40-μm cell strainer (Falcon, 352350 and 352340, respectively). The samples were incubated with 1 ml of red blood cell lysis buffer (15 mM NH_4_Cl, 12 mM NaHCO_3_, 0.1 mM EDTA, pH 8.0) for 2 min on ice. Leibovitz with 10% FBS and 1 mM EDTA was used for resuspension and washing for magnetic-activated cell sorting (MACS).

For pulmonary epithelial cell isolation, cells were resuspended in 400 μl of buffer with 30 μl of CD31 MicroBeads (130-097-418; Miltenyi Biotec, Bergisch Gladbach, Germany), 30 μl of CD45 MicroBeads (130-052-301; Miltenyi Biotec), and 30 μl of anti-Ter-119 MicroBeads (130-049-901; Miltenyi Biotec) and incubated for 30 min at 4 °C, followed by negative selection according to the manufacturer’s instructions. Cells were then resuspended with 400 μl of buffer with 30 μl of CD326 (EpCAM) MicroBeads (130-105-958; Miltenyi Biotec) and incubated for 30 min at 4 °C, followed by positive selection according to the manufacturer’s instructions. Isolated lung epithelial cells were used for the lung organoid culture.

For lung endothelial cell (LuEC) isolation, cells were resuspended in 400 μl of buffer with 30 μl of CD31 MicroBeads and incubated for 30 min at 4 °C, followed by positive selection according to the manufacturer’s instructions. Isolated LuECs were cultured with EC growth media (DMEM; Corning; MT10013CV, 20% FBS, 1’ glu-pen-strep; Gibco, USA; 10378016, 100 μg/ml endothelial cell growth factor (ECGS); Sigma; E2759, 100 μg/ml heparin; Sigma; H3149, 25 mM HEPES) on 0.1% gelatin (Sigma, G1393)-coated plates. Cultured LuECs were then isolated with CD31 MicroBeads and expanded until passage 3. Expanded LuECs were cryopreserved for lung organoid culture.

### Lung organoids

Lung epithelial cells (Ter119^-^/Cd31^-^/Cd45^-^/Epcam^+^) isolated from 7-10-week-old *Cracd* WT mice or *Cracd* KO were cultured with lung stromal cells in a 3D organoid air-liquid interface, as described previously^24, 45^. In brief, freshly sorted lung epithelial cells were resuspended in 3D organoid media (Dulbecco’s modified Eagle’s medium [DMEM]/F12 [Gibco, USA]), 10% FBS [Thermo Fisher Scientific], 1’ penicillin-streptomycin-glutamine [Thermo Fisher Scientific], and 1’ insulin-transferrin-selenium [Thermo Fisher Scientific.]) and mixed with LuECs at a ratio of 1:1. Cells containing 3D media were mixed with growth factor-reduced Matrigel (BD Biosciences) at a ratio of 1:1. The 100 ml of mixtures containing lung epithelial cells (5 × 10^3^) and LuECs (5 × 10^4^) were placed in the transwell insert (0.4-mm pore, Corning, Lowell, MA). After incubation for 30 mins at 37°C in an incubator, 500 ml of 3D media was placed in the bottom chamber to generate the liquid-air interface. Media were exchanged every other day.

### Mammalian cell culture

Human embryonic kidney 293T (HEK293T) and A549 cells were purchased from American Type Culture Collection (ATCC). The murine KP-1 cells were previously described ^23^. HEK293T cells were maintained in a DMEM medium containing 10% fetal bovine serum and 1% penicillin and streptomycin. A549 cells were maintained in Roswell Park Memorial Institute (RPMI) 1640 medium containing 10% fetal bovine serum and 1% penicillin and streptomycin. Cells were cultured at 37°C in a humidified incubator supplied with 5% CO_2_ air. Mycoplasma contamination was examined using the MycoAlert mycoplasma detection kit (Lonza).

## METHOD DETAILS

### qRT-PCR

RNAs were extracted by TRIzol (Invitrogen) and used to synthesize cDNAs using the iScript cDNA synthesis kit (Biorad). qRT-PCR was performed using an Applied Biosystems 7500 Real-Time PCR machine with the primers Target gene expression was normalized to that of mouse *Hprt1* and human *HPRT1*. Comparative 2^−ΔΔCt^ methods were used to quantify qRT-PCR results. (see Table S1 for primer information).

### Histology

#### Lung tissue

Lung tissues were perfused with cold PBS pH 7.4 into the right ventricle, fixed with 10% formalin, embedded in paraffin, and sectioned at 5-μm thickness. For H&E staining, sections were incubated in hematoxylin for 3-5 min and eosin Y for 20-40 s. For the immunohistochemistry analysis, sections were immunostained according to standard protocols^25^. For antigen retrieval, sections were subjected to heat-induced epitope retrieval pre-treatment at 120 °C using citrate-based antigen unmasking solution (Vector Laboratories, Burlingame, CA, USA). For immunofluorescence, after blocking with 10% goat serum in PBS for 30 min at ambient temperature, sections were incubated with primary antibodies (MKI67 [1:200], KRT19 [1:200], SYP [1:200], CGRP [1:200], CHGA [1:200], CDH1 [1:200], and ACTB [1:200]) overnight at 4 °C and secondary antibody (1:200) for 1 hr at ambient temperature. Sections were mounted with ProLong Gold antifade reagent with DAPI (Invitrogen). For chemically immuno-staining, sections were incubated with primary antibodies (CGRP [1:200], CHGA [1:200], ASCL1 [1:200], and NEUROD1 [1:200]) overnight at 4 °C and secondary antibody (1:200) for 1 hr at ambient temperature. 3,3’Diaminobenzidine (DAB) (Vector Laboratory) was used as the chromogens. Then, sections were dehydrated and were mounted with Permount (Thermo Fisher Scientific). Images were captured with the fluorescence microscope (Zeiss; AxioVision). See key resource table for antibody information.

#### Lung organoids (LOs)

LOs were harvested in ice-cold PBS. Then Matrigel was removed using cell recovery solution (Corning, Lowell, MA) for 1 hr at 4°C. Collected LOs were washed with ice-cold PBS two times, fixed with 10% formalin, embedded in paraffin, and sectioned at 5-μm thickness. For H&E staining, sections were incubated in hematoxylin for 3-5 min and eosin Y for 20-40 s. For the immunohistochemistry analysis, sections were immunostained according to standard protocols^25^. For antigen retrieval, sections were subjected to heat-induced epitope retrieval pre-treatment at 120 °C using citrate-based antigen unmasking solution (Vector Laboratories, Burlingame, CA, USA). For immunofluorescence, after blocking with 10% goat serum in PBS for 30 min at ambient temperature, sections were incubated with primary antibodies (CGRP [1:200], CHGA [1:200], HOPX [1:100], SPC [1:200], SCGB1A1 [1:200], and Ac-Tub [1:200]) overnight at 4 °C and secondary antibody (1:200) for 1 hr at ambient temperature. Sections were mounted with ProLong Gold antifade reagent with DAPI (Invitrogen). Images were captured with the fluorescence microscope (Zeiss; AxioVision). See key resource table for antibody information.

#### Cell lines

Cells were fixed for 20 min in 4% paraformaldehyde and permeabilized with 0.1% Triton X-100 (in PBS) for 10 min. After three PBS washes, cells were blocked with 2% bovine serum albumin (BSA) for 30 min at ambient temperature. Cells were then incubated with antibodies diluted in 2% BSA at 4°C overnight. After three PBS washes, the cells were incubated with phalloidin (Invitrogen) by shaking at ambient temperature in the dark for 1 h. Cells were washed three times with PBS in the dark and mounted in Prolong Gold Antifade Reagent (Invitrogen).

#### Microscopy

Immunofluorescent staining was observed and analyzed using a fluorescent microscope (ZEISS) and ZEN software (ZEISS).

#### Analyzing tumor heterogeneity index

Tumor heterogeneity was calculated based on the histomorphology of H&E staining. Each unique histomorphology in one tumor burden was scored as tumor heterogeneity index (fig. S2)

#### Virus production and transduction

Lentiviruses were produced using the 2^nd^-generation packaging vectors in 293T cells. 293T cells were cultured until 70%-80% confluent, and the media were replaced with antibiotics-free DMEM (10% FBS). After 1 hr of media exchange, cells were transfected with vector mixtures in Opti-MEM (Gibco, USA). To generate a vector mixture, pMD2.G (1.3 pmol), psPAX2 (0.72 pmol), DNA (1.64 pmol), and polyethyleneimine (PEI, 39 mg) were added to 800 ml of Opti-MEM and incubated for 15 mins. After 12 hrs of transfection, the media were exchanged with complete media (DMEM, 10% FBS, and 1× penicillin-streptomycin). The virus supernatant was collected after 24 hrs and 48 hrs and filtered with a 0.45-mm syringe filter (Thermo Fisher, CA, USA). pLenti-shCtrl (negative silencing control; Dharmacon), pLenti-shCRACD (Dharmacon; V3LHS_367334), and pLenti-shCracd (Dharmacon; V2LMM_57028) vectors were used for lentivirus generation. A549 and KP-1 cells were transduced by lentivirus containing shCtrl (control), or shCRACD or shCracd, respectively, with polybrene (8 μg/ml). Infected cells were selected using puromycin (2 μg/ml; Sigma). Adenovirus containing Ad-Cre, Ad-Cre-sgLacZ, and Ad-Cre-sgCracd vector were generated by Gene Vector Core at BCM. see Table S1 for shRNA and sgRNA sequences.

### scRNA-seq library preparation

#### Tissue preparation

Whole lungs were harvested from euthanized mice (Cracd WT or Cracd KO) after perfusing 10 ml of cold phosphate-buffered saline (PBS) into the right ventricle. The lung was digested in Leibovitz’s medium (Invitrogen) with 2 mg/mL Collagenase Type I (Worthington), 2 mg/mL Elastase (Worthington), and 2 mg/mL DNase I (Worthington) at 37 °C for 45 min. The tissue was triturated with a pipet every 15 min of digestion until homogenous. The digestion was stopped with FBS (Invitrogen) to a final concentration of 20%. The cells were filtered with a 70 μm cell strainer (Falcon) and spun down at 5,000 r/min for 1 min. The cell pellet was resuspended in red blood cell lysing buffer (Sigma) for 3 min, spun down at 5,000 r/min for 1 min, and washed with 1 mL ice-cold Leibovitz’s medium with 10% FBS. In single-cell RNA sequencing (scRNA-seq), digested lung cells were resuspended in 400 μl of buffer with 5 μl of anti-CD31-FITC (BD Biosciences, CA, USA), 5 μl of anti-CD45-APC (BD Biosciences), and 5 μl of anti-CD326 (EpCAM)-PE-Cy7 (Biolegend) and incubated for 30 min at 4 °C. Cells were then washed twice, followed by sorting of the epithelial cells (EpCAM+ / CD31- / CD45-) by fluorescence-activated cell sorting at the Cytometry and Cell Sorting Core at the Baylor College of Medicine.

#### Library

Single-cell Gene Expression Library was prepared according to the guideline for the Chromium Single Cell Gene Expression 3v3.1 kit (10× Genomics). Briefly, single cells, reverse transcription (RT) reagents, Gel Beads containing barcoded oligonucleotides, and oil were loaded on a Chromium controller (10× Genomics) to generate single-cell GEMS (Gel Beads-In-Emulsions), where full-length cDNA was synthesized and barcoded for each single cell. Subsequently, the GEMS were broken and cDNAs from each single cell were pooled, followed by cleanup using Dynabeads MyOne Silane Beads and cDNA amplification by PCR. The amplified product was then fragmented to optimal size before end-repair, A-tailing, and adaptor ligation. The final library was generated by amplification. The library was performed at the Single Cell Genomics Core at the Baylor College of Medicine.

### scRNA-seq data analysis

#### Data processing, clustering, and annotation

The Cell Ranger was used for demultiplexing, barcoded processing, and gene counting. The loom files were generated using the velocyto package ^46^. The R package Seurat^47^ and Python package Scanpy^48^ were used for pre-processing and clustering of scRNA-seq data with the loom files. UMAP was used for dimensional reduction, and cells were clustered in Seurat or Scanpy. Datasets were pre-processed, normalized separately. Each dataset was normalized separately and clustered by the “Leiden” algorithm ^49^. *Cracd* WT and *Cracd* KO datasets were combined using “ingest” function in Scanpy. Scanpy was used to concatenate the *Cracd* WT vs. KO dataset. Cells with more than 7000 counts reads were removed. Gene expression for each cell was normalized and log-transformed. The percentages of mitochondrial reads were regressed before scaling the data. Dimensionality reduction and Leiden clustering (resolution 0.5 ∼ 1) was carried out, and cell lineages were annotated based on algorithmically defined marker gene expression for each cluster (sc.tl.rank_genes_groups, method=‘wilcoxon’). Each cluster-specific gene list is shown in Table S2.

#### Gene set enrichment analysis (GESA)

AT2 cell clusters were isolated and then the DEGs between *Cracd* KO vs. *Cracd* WT in the AT2 clusters were identified by the Wilcoxon sum test and AUROC statistics using the Presto package v. 1.0.0. They were then subjected to GSEA using the fgsea package v. 1.16.0. The curated gene sets (C5) in the Molecular Signature Database (MsigDB) v. 7.5.1 were used for the GSEA using the msigdbr package.

#### Pathway score analysis

Scanpy with the ‘scanpy.tl.score_genes’ function or Seurat with the ‘AddModuleScore’ function were used for the pathway score analysis. The analysis was performed with default parameters and the reference genes from the gene ontology biological process or the Kyoto Encyclopedia of Genes and Genomes database ^50, 51^. The gene list for the score analysis is shown in Table S3.

#### Developmental state analysis

CytoTRACE (v. 0.3.3) ^27^ was used to predict the relative differentiation state of a single cell. The cells were given a CytoTRACE score according to their differentiation potential, with a higher score indicating higher stemness/fewer differential characteristics.

#### Human scRNA-seq data analysis

The public large cohort of scRNA-seq data sets (29 datasets; 556 samples; https://doi.org/10.5281/zenodo.6411867) were downloaded and analyzed ^29^. We analyzed only epithelial cell compartments (90,243 cells; 342 samples; 236 patients). The clusters were refined based on the neuroendocrine marker genes. For GSEA analysis, LUAD, LUAD mitotic, LUAD EMT, LUAD MSLN clusters were combined into as name of LUAD non-NE, and then GSEA of LUAD NE1 vs LUAD non-NE were analyzed described above.

## QUANTIFICATION AND STATISTICAL ANALYSIS

GraphPad Prism 9.4 (Dogmatics) was used for statistical analyses. The Student’s *t*-test was used to compare two samples. *P* values < 0.05 were considered statistically significant. Error bars indicate the standard deviation (s.d.) otherwise described in Figure legends. All experiments were performed three or more times independently under identical or similar conditions.

## Supplementary information

### Supplementary Figures

**Figure S1.**
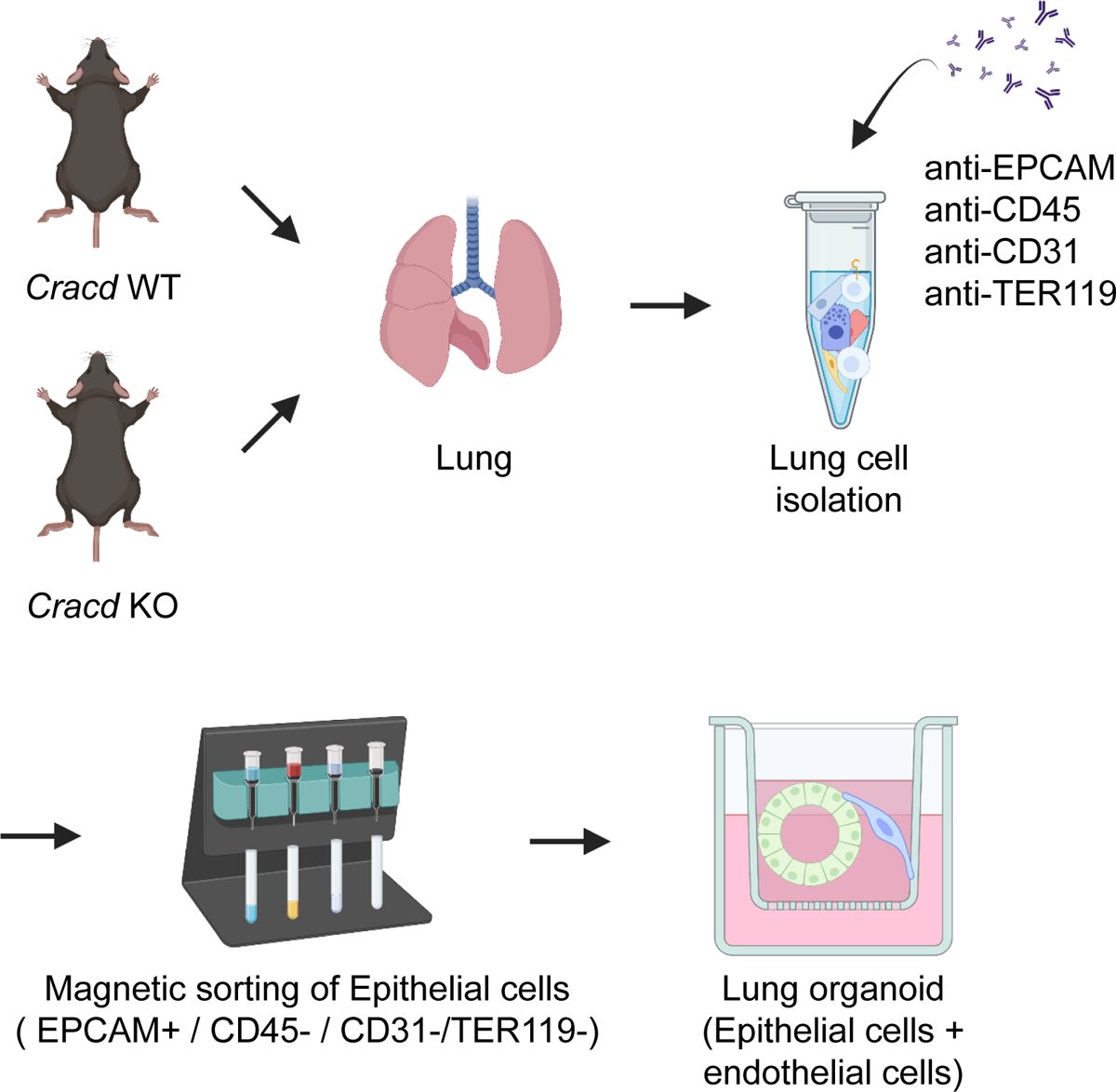
Illustration of the experimental scheme of lung organoid culture. Experimental scheme of LO culture. The lung epithelial cells were isolated from *Cracd* WT or *Cracd* KO murine lungs by magnetic-activated cell sorting (MACS). The lung epithelial cells (Ter119-/Cd31-/Cd45-/Epcam+) were co-cultured with lung endothelial cells (Cd31+) at a liquid-air interface to generate LOs.

**Figure S2.**
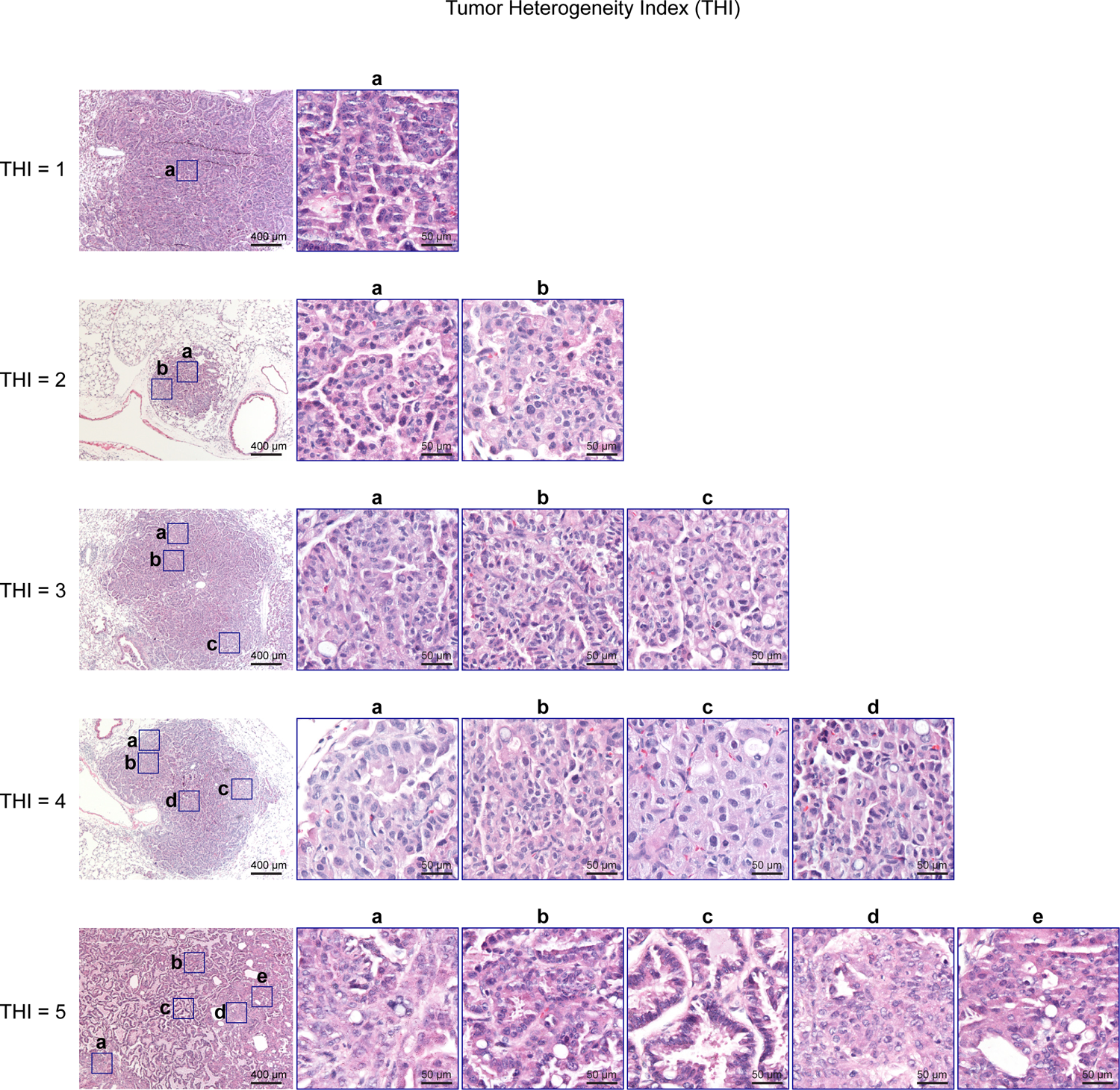
Evaluation of intratumoral heterogeneity by tumor heterogeneity index (THI). Intratumoral heterogeneity was assessed by calculating THI of each tumor. THI was determined based on the number of histologically different tumor types assessed by histomorphology of H&E staining.

**Figure S3.**
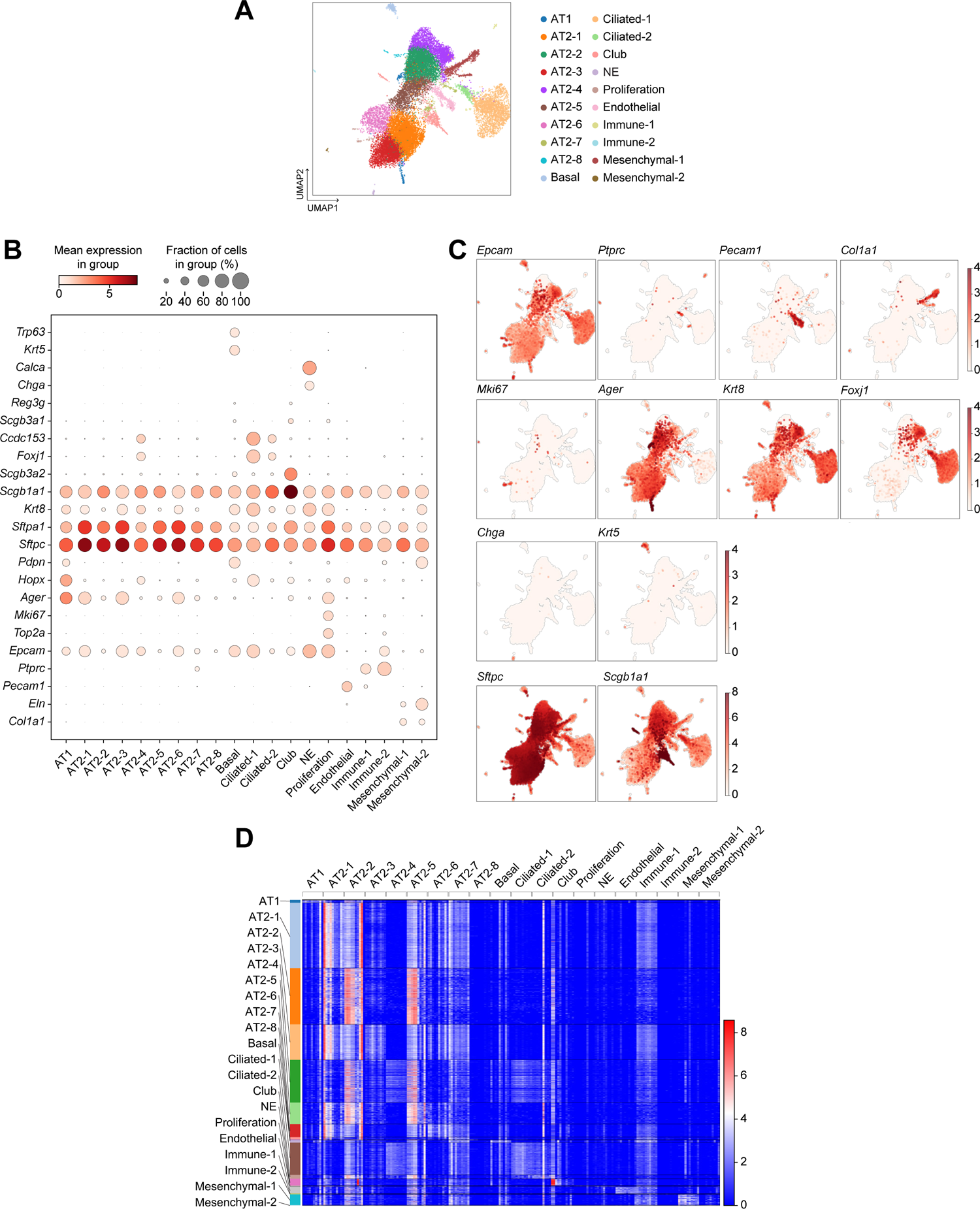
scRNA-seq of the murine pulmonary epithelial cells isolated from mice (*Cracd* WT vs. KO). **A,** UMAP of each cell cluster annotated by cell types. **B,** Dot plots depicting the expression of indicated lung epithelial cell type marker genes. **C,** Feature plots showing the expression of indicated lung epithelial cell type marker genes. **D,** Heatmap of each cluster-specific genes.

**Figure S4.**
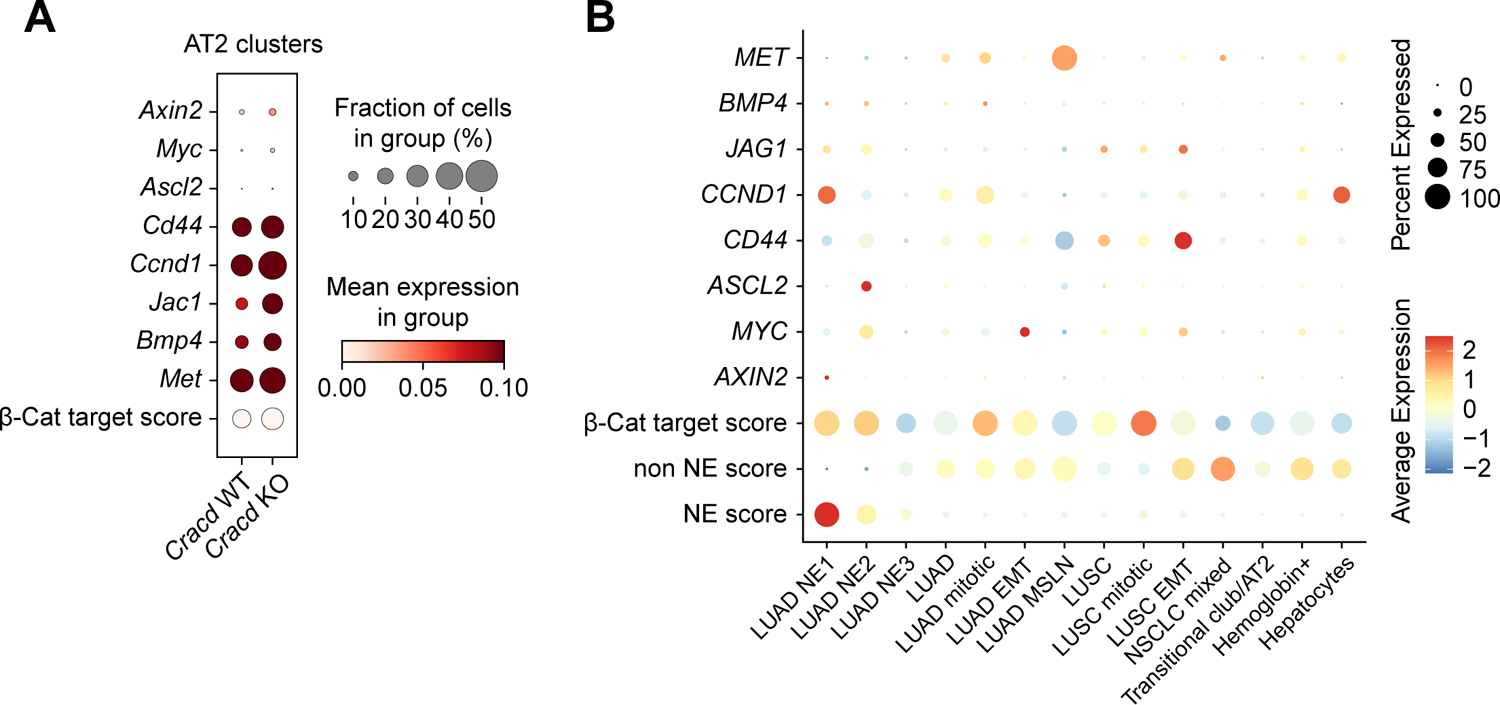
Analysis of WNT signaling activity. **A,** Dot plots depicting the expression of indicated genes and module score of β-Catenin target genes in AT2 cell clusters shown in Figure 4A. **B,** Dot plots depicting the expression of indicated genes and module score of β-Catenin target genes in scRNA-seq data shown in Figure 4H.

### Supplementary Tables

Table S1. Sequence information of primers and gRNA.

Table S2. Cluster specific gene list of scRNA-seq data.

Table S3. List of genes of each gene sets for moudule score analysis.

